# Rumen Bacteria and Serum Metabolites Predictive of Feed Efficiency Phenotypes in Beef Cattle

**DOI:** 10.1101/701300

**Authors:** Brooke A Clemmons, Cameron Martino, Joshua B Powers, Shawn R Campagna, Brynn H Voy, Dallas R. Donohoe, James Gaffney, Mallory M Embree, Phillip R Myer

## Abstract

The rumen microbiome is critical in ruminant nutrition and contributes to nutrient utilization and feed efficiency in cattle. Therefore, the objective of this study was to interrogate microbial and biochemical factors affecting divergences in feed efficiency in Angus steers using 16S amplicon sequencing and untargeted metabolomics. Average residual feed intake (RFI) was calculated, and steers were divided into low- and high-RFI groups. Features were ranked in relation to RFI through supervised machine learning on microbial and metabolite compositions. Residual feed intake was associated with several attributes of the rumen bacterial community. Low-RFI steers were associated with decreased bacterial α- (P=0.03) and β-diversity (P<0.001). Several serum metabolites were associated with RFI. Based on fold change (high/low RFI), low-RFI steers had greater abundances of pantothenate (P=0.02). Machine learning on RFI was predictive of both serum metabolomic signature and rumen bacterial composition (AUC ≥0.7). Log-ratio proportions of the bacterial classes Flavobacteriia over Fusobacteriia were enriched in low-RFI steers (F=6.8, P=.01). Greater proportions of pantothenate-producing bacteria, such as Flavobacteriia, and/or reductions in Fusobacteriia may result in improved nutrient utilization in low-RFI steers. Pantothenate and Flavobacteriia may serve as potentially novel biomarkers to assess or predict feed efficiency in Angus steers.

## Introduction

The United States is the largest producer of beef, and the beef industry accounts for a retail equivalent of $105 billion ^1^. Over the next decade, demand for US exportation of beef is expected to increase ^2^. Given the rapid reduction in natural resources and substantial growth of the human population expected in the coming decades, it is imperative to develop novel agricultural approaches in order to increase the global food supply with limited resources ^3^. Ruminants, including beef cattle, rely on the fermentation of feedstuffs to provide energy for the animal. The rumen microbiome in cattle is fundamental for the successful conversion of plant matter to energy substrates for the animal via fermentation ^4^. This microbiome also supplies the host animal with other important nutrients such as vitamins and protein ^4^. Identifying and exploiting factors that affect the efficiency of this conversion in beef cattle will result in increased animal protein supply without increasing input resources.

Rumen microbes produce metabolites that are released into the rumen lumen and can be absorbed through the rumen epithelium or through the epithelium in the lower gastrointestinal tract ^4^. The rumen microbes are responsible for the production of approximately 70% of the energy supply to the ruminant, including production of organic acids such as acetate and propionate ^5^. Differences in the production of these metabolites, as well as variation in rate and quantity of absorption, can contribute to variation in nutrient utilization and efficiency of the ruminants, and may lead to physiological or phenotypic changes ^6,7^. However, it can be difficult to distinguish the origin of many metabolites between those of endogenous origin and metabolites of microbial origin. Although associations between the rumen microbiome and physiological changes in the host have been identified ^8,9^, the mechanisms driving these changes are still unknown and whether foundational, or keystone, species are responsible for the divergences in feed efficiency and other phenotypes.

In order to address these critical knowledge gaps, we used a combination of microbial genomics, metabolomics, and bioinformatics to further define variations in feed efficiency as determined by the divergence in residual feed intake (RFI). Determination of the complex associations and networks between the rumen microbiome, host metabolome, and differences in host phenotype can be facilitated by novel utilization of bioinformatics and machine learning to discover physiological patterns and microbial factors. By taking a multidisciplinary approach, research can move beyond correlation to identify variables accounting for differences in host phenotype.

The objective of this study was to identify the microbial and biochemical biomarkers mediating variation in feed efficiency in cattle. To accomplish this, we analyzed the relationships among RFI, the rumen bacterial community, and the serum metabolome.

## Results

### Sequencing information

A total number of 50 samples underwent microbial DNA extraction. Bacterial community composition was determined by amplifying and sequencing the V1-V3 hypervariable region of the 16S rRNA gene. 21,734,148 number of sequences were present following quality control and chimera removal. An average of 48,048 ± 41,628 sequences was present in each sample.

### Bacterial community diversity

After binning reads at 97% similarity, a total of 21,401 OTU were detected. Alpha-diversity was measured by equitability, Simpson’s Evenness, observed OTU, Good’s coverage, chao1, and Shannon’s Diversity Index. Alpha-diversity metrics did not differ between low- and high-RFI steers at the end of the study (Table 1), including equitability (P=0.24; Table 1), Simpson’s Evenness (P=0.19; Table 1), Observed OTU (P=0.78; Table 1), Good’s coverage (P=0.14; Table 1), chao1 (P=0.78; Table 1), and Shannon’s Diversity Index (P=0.07; Table 1). Beta diversity of the rumen bacterial communities also changed significantly over time (PERMANOVA: F = 422, P< .001, Figure 1) with highly ranked bacterial classes Gammaproteobacteria, Alphaproteobacteria, and Bacteroidia diverging in time.

**Table 1.**
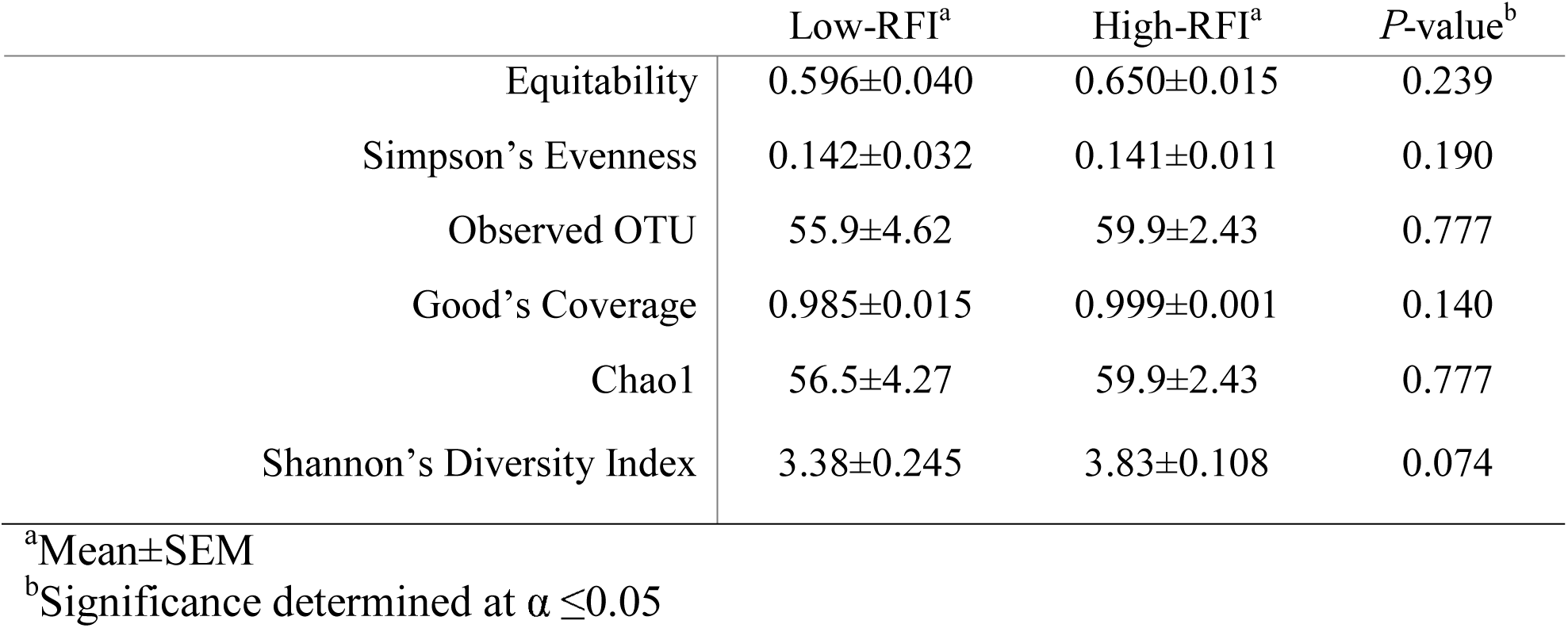
Sequence and alpha-diversity statistics of the 16S rRNA gene sequences for bacterial populations in low- and high-RFI steers.

**Figure 1.**
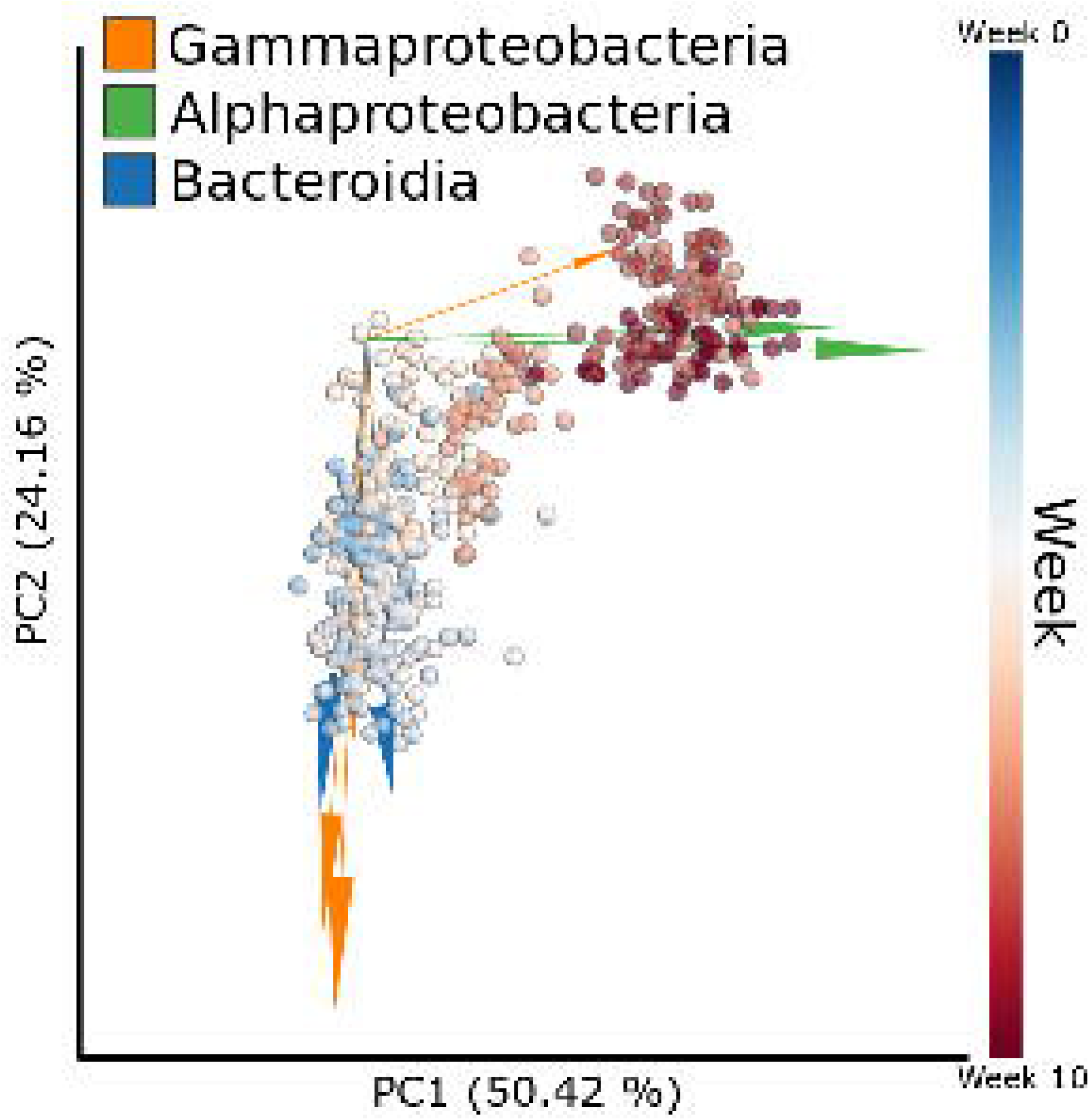
Compositional biplot beta diversity generated through Robust Aitchison PCA comparing microbial compositions over time (weeks) with arrows representing the highly ranked bacterial features colored by class level taxonomy.

### Biochemical and microbial predictors of RFI

A total of 114 known metabolites were identified. Residual feed intake was predictive of serum metabolomic composition (AUC=.75, Figure 2A) and rumen bacterial composition (AUC=.74, Figure 2B) at week 10 using LSVC and RF machine learning respectively. Many serum metabolites were identified as predictive of high- and low-RFI between steers, one such metabolite being pantothenate. (Supplementary Table 1). The serum pantothenate proportion was also found to be significantly different between low- and high-RFI steers (F=5.89, P=0.02, Figure 3A). Additionally, among the highly ranked microbial classes the rumen bacterial classes of Flavobacteriia and Fusobacteriia (Supplemental Table 2). The log-ratio of Flavobacteriia (numerator) and Fusobacteriia (denominator) was significantly increased in low-RFI steers (F=6.8, P=0.01) (Figure 3B). Furthermore, when using Cyanobacteria, a class found at low but constant abundance across all samples as the denominator. The log ratio of Flavobacteriia (numerator) and Cyanobacteria (denominator) correlated well with pantothenate abundance (Figure 3C).

**Figure 2.**
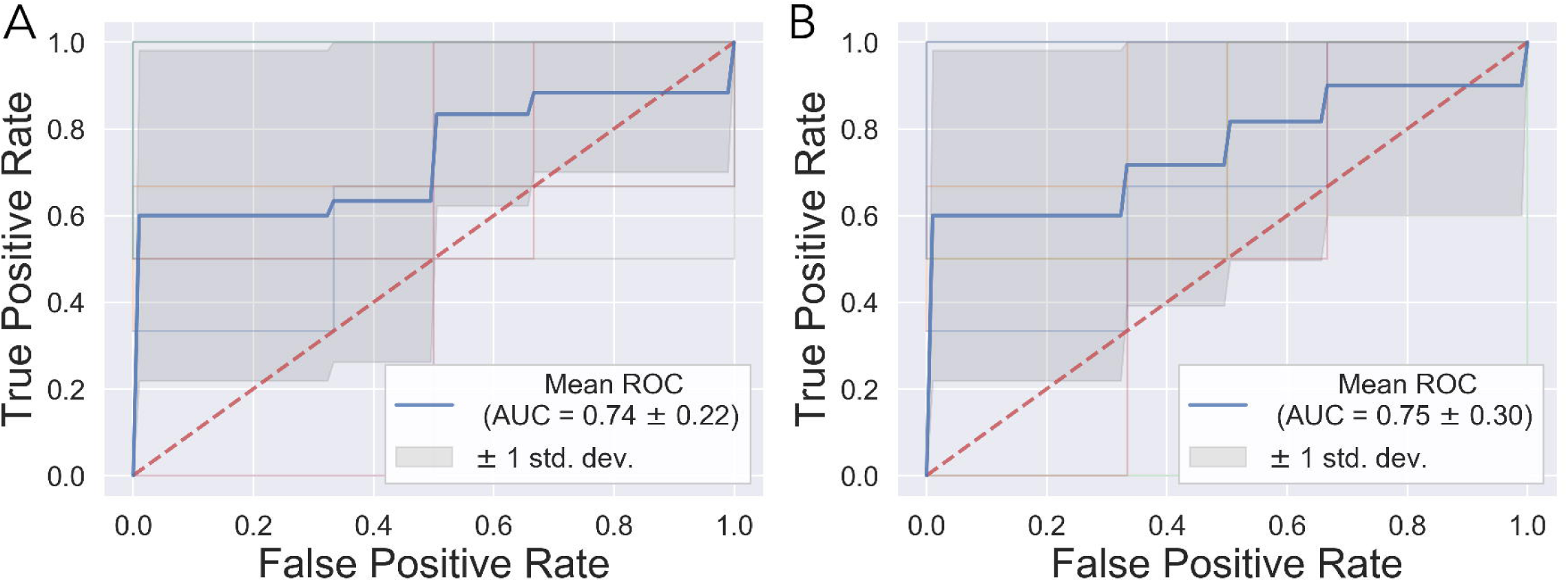
Ten-fold stratified K-Folds cross-validation ROC curves for the prediction accuracy (AUC) for bacterial (A) and metabolite (B) compositions of RFI.

**Figure 3.**
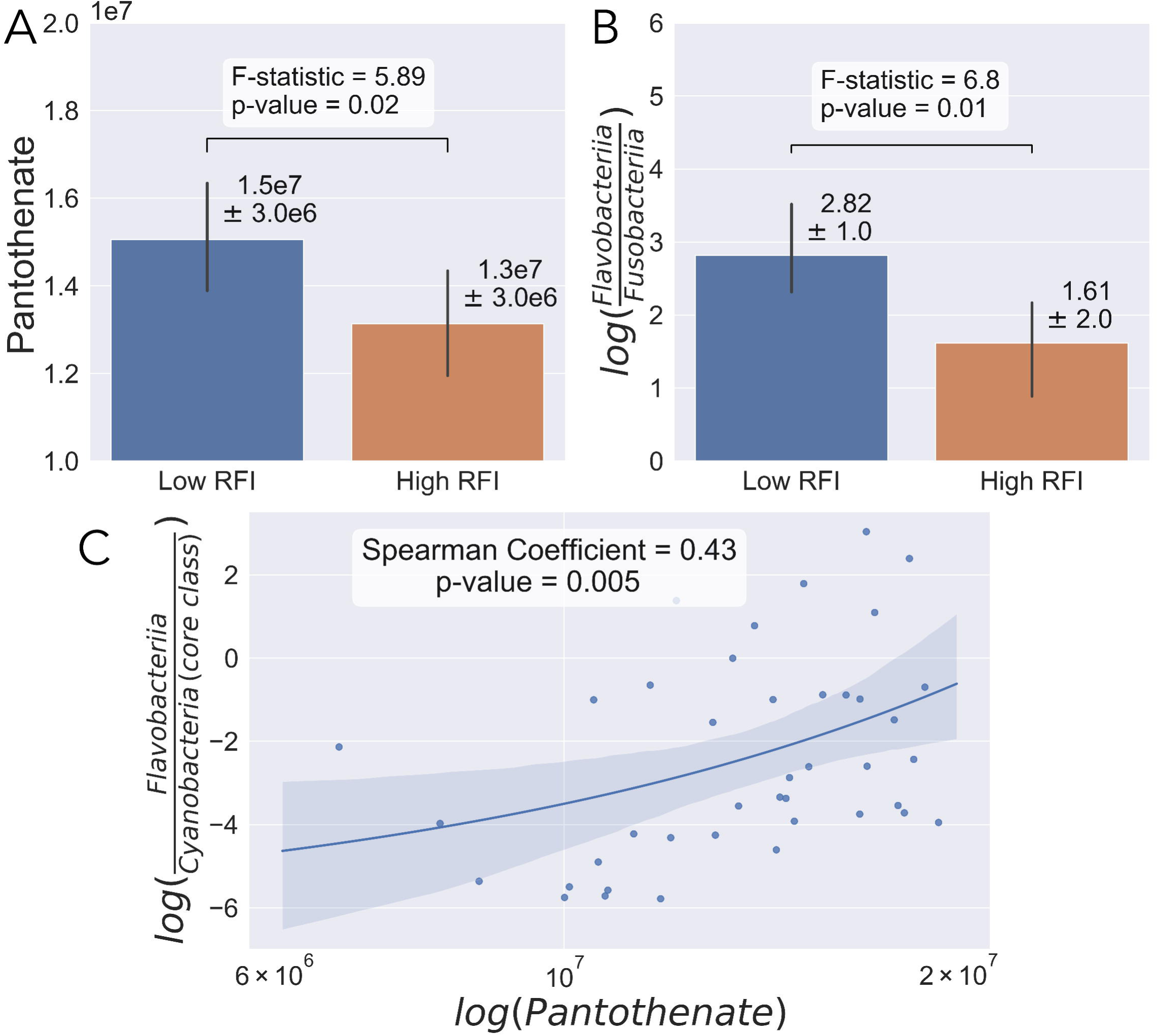
Analysis of week ten microbial compositions and metabolite abundances. Pantothenate abundance compared between low- and high-RFI (A). Log-ratio proportions of highly ranked microbes Flavobacteriia (numerator) and Fusobacteria (denominator) compared between low- and high-RFI (B). Regression plot between the log-ratio of Flavobacteriia (numerator) / Cyanobacteria (denominator) and the log-scaled pantothenate abundance.

## Discussion

Total beef consumption in the United States is greater than 20 billion pounds annually, and is the most consumed red meat product in the United States ^1^; however, declining land resources and increasing human populations place pressure on producers to improve production efficiency. Therefore, researchers and producers are charged with finding novel approaches and methods for reducing inputs while increasing animal protein supply to meet the needs of an expected global population exceeding 9 billion people by the year 2050 ^3^. Given this need, targeting phenotypes that improve efficiencies, such a feed and reproductive efficiencies, or reduce negative environmental impacts, including methane production and excess nitrogen release, will ultimately improve beef and livestock agriculture on a global scale. The rumen microbiome contributes significantly to the breakdown of low-quality feedstuffs, such as forages, and may be responsible for much of the variation observed in ruminant feed efficiency phenotypes (Bergman 1990). Understanding the relationships between host phenotypes and the rumen microbiome may provide novel methods for improving feed efficiency in ruminants.

In this study, measurements of α-diversity did not differ between the two groups. In other ecosystems, increased biodiversity is associated with greater success and resilience of the ecosystem ^10,11^; however, the data presented in this study suggest that α-diversity may not be a significant contributing factor to feed efficiency phenotypes in stable bacterial communities in growing beef steers. Previous studies have also observed the same relationships between rumen bacterial α-diversity and feed efficiency phenotypes in steers ^12,13^. Other factors, such as divergences in individual taxa or functionality of the rumen microbiota, may play a greater role in dictation of host feed efficiency phenotypes.

Given the relationship between feed efficiency phenotypes and the rumen bacterial communities, it may be possible to identify specific rumen microbes and serum metabolites associated with RFI. In the human vagina, for example, *Lactobacillus spp.* is often associated with pregnancy success and woman reproductive tract health ^14-16^. In the rumen, specific taxa, even if at lower abundances, may cause distinct variation in feed efficiency phenotypes ^17^. The lack of differences in α-diversity in previous studies suggests that divergences in feed efficiency phenotypes may be the result of dissimilarities at a finer resolution, such as individual taxa and metabolites, rather than global changes in the microbial communities and cumulative metabolites. Differences in serum and rumen metabolites may provide indications of feed efficiency and could be developed into a method for on-farm detection of feed efficiency.

The rumen microbiome produces several vital nutrients for the host animals, including organic acids that serve as glucogenic precursors, as well as proteins and vitamins ^4^. A nutrient produced by the rumen microbiota is pantothenate. Pantothenate plays a significant role in the metabolism of fatty acids in ruminants and other species ^18,19^. In this study, pantothenate was identified as a potential biomarker of RFI and was shown to be significantly enriched in low-RFI compared to high-RFI steers. Pantothenate is a key component of coenzyme A (CoA), which is required to perform a variety of functions in intermediary metabolism of ruminants ^20^. Namely, CoA is responsible for the transfer of fatty acid components into and out of the mitochondria ^21^. Pantothenate is produced by several species of bacteria in the rumen, and can then be released into the rumen lumen to be absorbed by the host animal. One class of bacteria that can generate pantothenate in the rumen are Flavobacteriia.

In this study, Flavobacteriia was also identified as a potential biomarker of RFI. However, because the absolute population of microbes in each sample is unknown, log-ratios were used to compare between low- and high-RFI ^22,23^. It was found in this study that the log-ratio of Flavobacteriia and Cyanobacteria were well correlated to pantothenate abundance. Cyanobacteria was identified as a class with low-ranked and non-fluctuating proportions across all samples and used as the denominator in the log-ratio. Although, Cyanobacteria are oxygenic phototrophic bacteria they are often found at low but constant proportions in the rumen ^24,25^ and thought to be possibly misclassified from the class Melainabacteria ^26^. By using the log-ratio of Flavobacteriia and Cyanobacteria, the assumption can be made that this correlation is caused by an enrichment in Flavobacteriia and not a decrease in Cyanobacteria. This information supports that more efficient steers are associated with increased proportions of both Flavobacteriia and pantothenate.

In contrast to the log-ratio used in the correlation, the log-ratio of Flavobacteriia and Fusobacteria do not allow the same assumptions to be made as used above. This suggests that low-RFI animals could be caused by an increase or decrease of Flavobacteriia or an increase or decrease in Fusobacteria. *Fusobacterium necrophorum* of the class Fusobacteria is a known opportunistic pathogen and causative agent of liver abscesses in cattle ^27^. Although *F. necrophorum* is a normal rumen inhabitant ^28^, it is known to be enriched in high-grain diets ^29^. This enrichment causes *F. necrophorum* to leak into portal circulation where it is then trapped in the liver causing abscesses ^30^. Fusobacteriia and Flavobacteriia both identified as prospective important biomarkers here may play joined roles in regulating feed efficiency. However, further experimentation is needed to delineate this relationship.

Pantothenate may indicate greater feed efficiency. The relationship between pantothenate and Flavobacteriia could provide insight beyond the mechanisms accounting for some variability in feed efficiency, by possibly serving as biochemical and microbial biomarkers in the serum and rumen, respectively. These biomarkers could allow producers to identify and select animals of greater feed efficiency. Metabolites and microbes predictive of efficiency phenotypes in cattle are not only imperative to partially explaining divergences in feed efficiency, but also to the selection of microbial communities related to efficient animals. These insights may also lead to the ability to select for an optimal rumen microbiome.

This study identified potential microbial and biochemical biomarkers that were used to determine extremes in feed efficiency in steers. Although notable correlations between pantothenate and feed efficiency were identified, linking, and perhaps predicting, the functional capacity of the rumen and its microbiome, specifically Flavobacteriia, through serum pantothenate offers the potential to use serum biochemistry as an indicator in identification of feed efficient cattle. Additionally, although it has yet to be determined to what degree the rumen microbiome influences the host, or the host influences the rumen microbiome, the present study identified several key physiological elements that may impact or predict microbial community structure (e.g. RFI), or predictive of RFI (i.e. the serum metabolome and rumen bacterial community). As producers and researchers alike search for sources of variation in feed efficiency in cattle with the intent to optimize cattle productivity, methods to predict feed efficiency, such as use of microbial and biochemical markers could ultimately be used to improve the selection for feed efficient cattle.

## Materials and Methods

This study was approved and carried out in accordance with the recommendations of the Institutional Animal Care and Use Committee at the University of Tennessee, Knoxville.

### Animal experimental design and sample collection

Fifty weaned steers of approximately 7 months of age were housed at the Plateau Research and Education Center in Crossville, TN ^31^. Animals weighed 264 ± 2.7 kg at the beginning of the study and transitioned to a backgrounding diet for 14 days prior to the start of the trial. Diet consisted of 80% corn silage, 10% cracked corn, and 10% protein supplement (11.57% crude protein and 76.93% total digestible nutrients with 28 mg monensin/kg on a dry matter basis). A 70-day feed efficiency trial was administered following the acclimation period. Steers were adapted to the GrowSafe^©^ system during that adaptation period. Body weight (BW) was measured at 7-day intervals and daily feed intake measured using the GrowSafe^©^ system for the length of the 70-day feed efficiency trial. Feed efficiency was determined using RFI ^32^. At the conclusion of the trial, steers were ranked based on RFI and samples from the low- and high-RFI animals were utilized for subsequent analyses. Low- (*n*=14) or high- (*n*=15) RFI was determined as 0.5 SD below or above the mean RFI, respectively.

Weekly, approximately 9mL of blood was sampled via venipuncture from the coccygeal vein into serum separator tubes (Corvac, Kendall Health Care, St. Louis, MO). Blood samples were centrifuged at 2,000x*g* for 20 min at 4°C. Serum was decanted into 5mL plastic culture tubes and stored at -80°C for further analyses. Approximately 100 mL of rumen content was collected via esophageal tubing and any content remaining on the filtered strainer was also collected ^33^. Samples were transferred to 50 mL conical tubes, pH was measured using a portable pH meter, and stored at -80°C until further processing.

### DNA extraction and amplification

Rumen samples were centrifuged at 4,000 rpm for 15 min, and the supernatant was decanted and discarded. Approximately 0.5 mL of the remaining pellet was aliquoted for DNA extraction using the PowerViral® Environmental RNA/DNA Isolation Kit (Mo Bio Laboratories, Inc., Carlsbad, CA, USA). The 16S rRNA gene was amplified using 27F ^34^ and 534R ^35^ modified for Illumina sequencing following standard protocols Q5® High-Fidelity DNA Polymerase (New England Biolabs, Inc., Ipswich, MA, USA). Following amplification, PCR products were verified with a 2% agarose gel electrophoresis and purified using AMPure XP bead (Beckman Coulter, Brea, CA, USA). The purified amplicon library was quantified and sequenced on the MiSeq Platform (Illumina, San Diego, CA, USA) according to standard protocols ^36^. Raw fastq reads were de-multiplexed on the MiSeq Platform (Illumina, San Diego, CA, USA).

### Phylogenetic analysis

All raw sequencing data were trimmed of adapter sequences and phred33 quality filtered at a cutoff of 20 using Trim Galore ^37^. All remaining sequences were then filtered for PhiX, low-complexity reads and cross-talk ^38^. 16S taxonomic sequence clustering and classification was performed with USEARCH UNOISE and SINTAX (v10.0.240) ^39,40^ with the RDP 16S rRNA database ^41^. Each sample was filtered for sequencing depth at a minimum of 2,000 reads per sample ^42^. Samples with fewer than 2,000 sequences were considered too low for adequate depth and were excluded from subsequent analyses.

### LC-MS analysis

Serum samples (50 μL) from each steer were extracted for metabolomic analysis using 0.1% formic acid in acetonitrile:water:methanol (2:2:1), as described previous ^43^. Metabolites were separated using a Synergy Hydro-RP column (100 ×2 mm, 2.5 μm particle size). Mobile phases consisted of A: 97:3 H2O:MeOH with 11 mM tributylamine and 15 mM acetic acid and B: MeOH. The gradient consisted of the following: 0.0 min, 0% B; 2.5 min 0% B; 5.0 min, 20% B; 7.5 min, 20% B; 13 min, 55% B; 15.5 min, 95% B; 18.5 min, 95% B; 19 min, 0% B, and 25 min, 0% B. Flow rate was set to a constant 0.200 mL/min and the column temperature remained at 25°C. The autosampler tray was maintained at 4°C and 10 μL of sample was injected into the Dionex UltiMate 3000 UPLC system (Thermo Fisher Scientific, Waltham, MA). Electrospray ionization was used to introduce the samples into an Exactive Plus Orbitrap MS (Thermo Fisher Scientific, Waltham, MA), using an established method ^43,44^.

Raw files obtained from Xcalibur MS software (Thermo Electron Corp., Waltham, MA) were converted into the mzML format using ProteoWizard ^45^. The converted files were imported into MAVEN (Metabolomic Analysis and Visualization Engine for LC-MS Data), a software package ^46^. Peaks for the known metabolites were picked in MAVEN, which automatically performs non-linear retention time correction and calculates peak areas across samples, using a preliminary mass error of ±20 ppm and retention time window of five min. The UTK Biological and Small Molecule Mass Spectrometry Core (BSMMSC) has replicated and expanded the method of Rabinowitz and coworkers ^44^ and final metabolite annotations were made using a library of 263 retention time-accurate m/z pairs taken from MS1 spectra. The annotation parameters have been verified previously with pure standards as part of establishing the method. For a metabolite to be annotated as a known compound, the eluted peak had to be found within two min of the expected retention time, and the metabolite mass had to be within ±5 ppm of the expected value. Metabolite identities were confirmed using the MAVEN software package ^46^, and peak areas for each compound were integrated using the Quan Browser function of the Xcalibur MS Software (Thermo Electron Corp., Waltham, MA).

### Statistical analyses

Beta diversity analysis was performed through Robust Aitchison PCA via deicode ^47^ and the resulting biplot was visualized through EMPeror ^48^. Compositional transformation of both the bacterial and metabolite data tables were performed through the centered log-ratio transform (clr) ^49^ with a pseudo count of one. Feature ranking and supervised machine learning was performed on clr transformed data through Random Forests ^50^. The clr transform and Permutational Multivariate Analysis of Variance (PERMANOVA) for beta diversity significance was performed through scikit-bio (http://scikit-bio.org/), data wrangling through pandas ^51^, visualization through seaborn ^52^ and matplotlib ^53^. Random Forest Classification (RF) was performed on bacterial compositions and Linear Support Vector Classification (LSVC) was performed on metabolite compositions with default parameters through scikit-learn ^54^. Additionally, ten-fold stratified K-Folds cross-validation was used to generate receiver operating characteristic (ROC) curves to evaluate the prediction accuracy under the curve (AUC) for each classification through scikit-learn.

Measurements of α-diversity, including equitability, Simpson’s Evenness E, Shannon’s Diversity Index, and Observed OTU, were assessed for normality using SAS 9.4 (SAS Institute, Cary, NC). All variables were found to follow a non-normal distribution, and were analyzed using Wilcoxon Rank Sum and Kruskal Wallis test.

## Data Availability

The datasets generated and/or analyzed during the current study are available from the corresponding author(s) on reasonable request.

## Acknowledgements

The authors thank Emily Melchior and the staff at the Plateau Research and Education Center in Crossville, TN for their technical assistance. This study was supported by Ascus Biosciences, Inc. (Grant No. A17-0146-003) and USDA-NIFA Hatch/Multistate Project W4177 - TEN00538 - Enhancing the Competitiveness and Value of U.S. Beef; Accession Number: 1016984.

## Author Contributions

BC contributed to study design, sample and data collection, data analysis, and preparation of manuscript. CM contributed to study design, data analysis, and preparation of manuscript. JP contributed to data analysis and preparation of manuscript. SC contributed to data analysis and preparation of manuscript. BV contributed to study design, data analysis, and preparation of manuscript. DD contributed to study design, and preparation of the manuscript. JG contributed to study design, and preparation of the manuscript. ME contributed to study design, data analysis, and preparation of manuscript. PM contributed to study design, sample and data collection, data analysis, and preparation of manuscript.

---

Supplementary Table 1. Bacterial class ranks in relation to RFI (High/Low) from Random Forest Classification feature rankings

Supplementary Table 2. Metabolite ranks in relation to RFI (High/Low) from Linear Support Vector Classification

